# Identification of a new family of “megaphages” that are abundant in the marine environment

**DOI:** 10.1101/2021.07.26.453748

**Authors:** Slawomir Michniewski, Branko Rihtman, Ryan Cook, Michael A. Jones, William H Wilson, David J Scanlan, Andrew Millard

## Abstract

Megaphages – bacteriophages harbouring extremely large genomes – have recently been found to be ubiquitous, being described from a variety of microbiomes ranging from the animal gut to soil and freshwater systems. However, no complete marine megaphage has been identified to date. Here, using both short and long read sequencing, we assembled >900 high-quality draft viral genomes from water in the English Channel. One of these genomes included a novel megaphage, Mar_Mega_1 at >650 Kb, making it one of the largest phage genomes assembled to date. Utilising phylogenetic and network approaches, we found this phage represents a new family of bacteriophages. Genomic analysis showed Mar_Mega_1 shares relatively few homologues with its closest relatives, but, as with other mega-phages Mar_Mega_1 contained a variety of auxiliary metabolic genes responsible for carbon metabolism and nucleotide biosynthesis, including isocitrate dehydrogenase [NADP] and nicotinamide-nucleotide amidohydrolase [PncC] which have not previously been identified in megaphages. The results of this study indicate that phages containing extremely large genomes can be found in abundance in the marine environment and augment host metabolism by mechanisms not previously described.

## Introduction

Phages – viruses that prey on bacteria – are the most abundant biological entities on Earth. Although they are ubiquitous and highly diverse components of the microbiome [1] the majority of known phages contain genomes smaller than 200 kb [2]. However, advances in the field of viral metagenomics led to the recent discovery of megaphages – phages with extremely large (>540 kb in length) genomes. The first identified megaphages, Lak phages, contained alternatively coded genomes and were present in both animal and human gut microbiota [3]. This was followed by the identification of multiple megaphage genomes, including the largest known phage genome 735 kb in length, from a range of viromes across Earth’s ecosystems [4]. Currently megaphages have been found in human and animal microbiomes [3, 5] soil and deep subsurface environments [4] and freshwater lakes [4, 6]. However, megaphages thus far have not been reported from marine systems.

### Results

Using a combination of Illumina and MinION sequencing, three marine viral communities isolated from waters of the western English Channel and Plymouth Sound were investigated (Supplementary Methods) resulting in the reconstruction of 23,179 putative viral contigs (acc:ERZ2485795), with 972 high-quality draft genomes of which 367 are predicted to be complete based on MIUViG standards [7] (Table S1). These included six phage genomes with lengths >200 kb and one exceptionally large, 656,628 bp genome, Mar_Mega_1 (acc:GCA_910591635). This genome comprised 1062 predicted genes including one tmRNA, 50 tRNAs and 1011 coding sequences (CDS). However, a combination of approaches including Blast, hmm searching and Phyre2 analyses resulted in the function being assigned for only 268 proteins (Table S2). The large genome size of Mar_Mega_1 makes it one of the largest phage genomes assembled to date and puts it in the range of “megaphages”. Comparison of Mar_Mega_1 with known phages showed no significant similarity at the nucleotide level. To determine its relationship with other phages a phylogeny was built using the amino-acid sequence of the terminase large subunit (Figure 1). All Lak-like megaphages formed a single clade, as was previously described [3], whilst Mar_Mega_1 formed a cluster with the largest megaphages (genome lengths >630 kb). Our newly identified phage is in a sister group to the uncultured phages LR756502 and LR745206 [4], isolated from a freshwater lake in France and sub-surface sample in Japan, respectively. However, the long branch lengths suggest that Mar_Mega_1 is only distantly related to these freshwater megaphages.

**Fig1.**
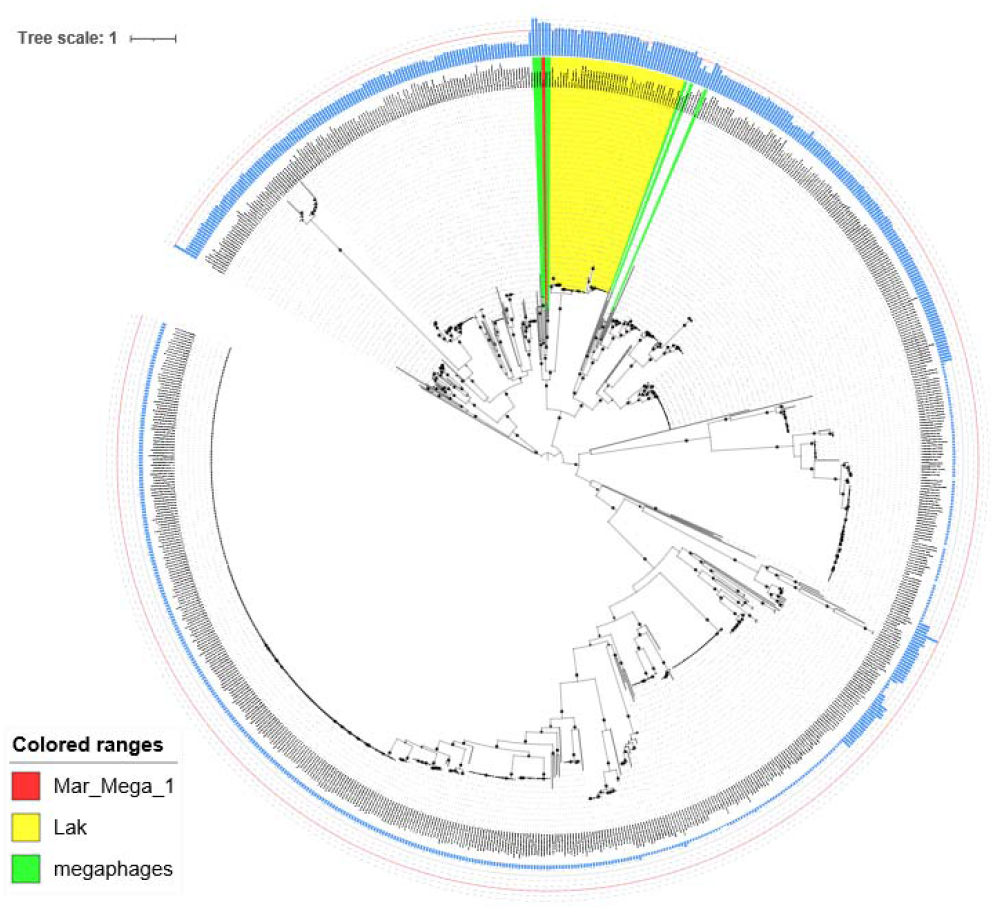
Phylogenetic analysis of selected phages and all megaphages (including incomplete megaphage genomes) based on the amino acid sequence of the terminase large subunit. Protein sequences were aligned with MAFFT and trees constructed with IQ-TREE, based on the LG+R7 model of evolution with 1000 bootstrap replicates. Mar_Mega_1 was marked with red, Lak phages with yellow and the remaining megaphages with green colour. Filled circles denote bootstrap values >70%, with the size of the circle proportional to the bootstrap value.

To further investigate the relationship of Mar_Mega_1 with its closest relatives, a core gene approach was used using parameters recommended for defining phage families [8]. All three phages share a core of only 125 genes that constitutes between 12.3% (Mar_Mega_1) and 13.2% (LR745206) of genes in each phage, suggesting these phages do not form a single family based on current definitions [8]. Phages LR756502 and LR745206 share nearly twice as many genes with each other (30.6 % - 31.5 %) than either phage does with Mar_Mega_1 (14.4 % - 17.5 %). Thus, suggesting Mar_Mega_1 represents a new family of marine megaphages based on current standards [8].

Having established Mar_Mega_1 as the first representative of a new family, we sought to establish its distribution in the marine environment. Although Mar_Mega_1 was present only in the samples taken from Plymouth Sound (Table S1), we have estimated that it is as abundant as cultivated phages that infect marine bacteria such as *Pelagibacter* and *Synechococcus* (e.g. *Synechococcus* phage S-SKS1, *Pelagibacter* phage HTVC115P and Lentibacter phage vB_LenP_ICBM2) through read mapping (Figure 2). However, the abundance of Mar_Mega_1 might be underestimated, as the majority of viral particles larger than 0.22 μm should have been removed during the filtration step. To examine the global distribution of Mar_Mega_1 we first tried using a read mapping approach against the TARA oceans virome dataset [9]. Despite several hundreds of reads mapping, no sample passed the accepted threshold of >1x coverage across 70% of the genome that is commonly used [10]. Therefore, we took an alternative approach using the TerL sequence from Mar_Mega_1 to query the TARA contigs via BlastP. Phylogenetic analysis of the top 100 phage hits revealed a further nine phages that are sister to the group containing Mar_Mega_1, suggesting related megaphages are present in the TARA oceans dataset [9]. As these proteins were found on genome fragments, it was not possible to compare total genome content with Mar_Mega_1. However, their placement within the same cluster, suggests they are further representatives of marine megaphages.

**Fig2.**
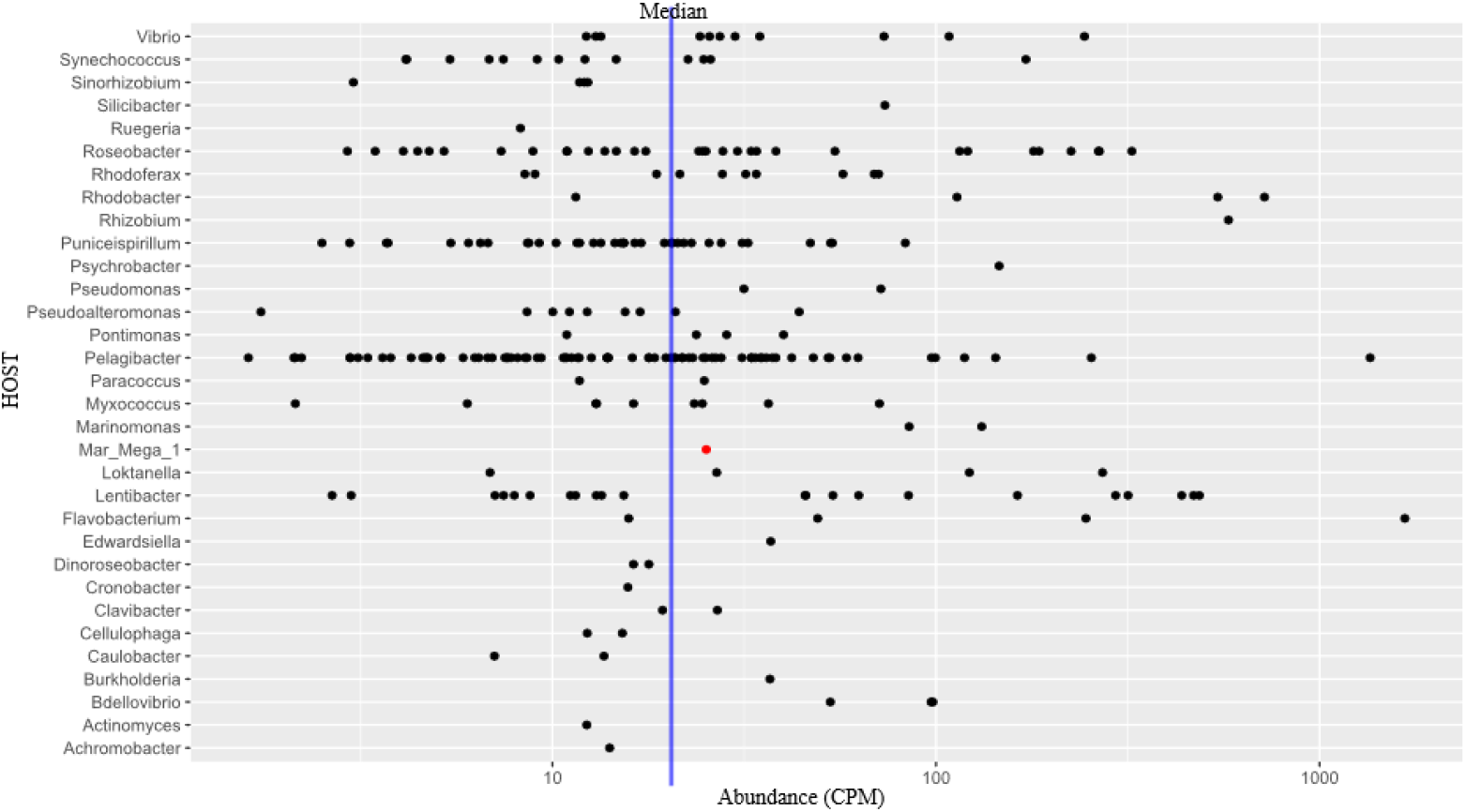
Abundance of viral contigs associated with known phages in the Plymouth Sound virome. Abundance is represented by counts per million. Each black dot represents a viral contig that could be associated with a known phage, based on mash similarity or clusters in the same group as known phages with vContact2. Mar_Mega_1 is identifiable by a red dot. The median coverage for all contigs that could be associated with known phages is identified by the blue line. Host information was extracted from known phages.

A number of methods were tested to determine a putative host for megaphage Mar_Mega_1 including CRISPR-spacer database analysis (see Supplementary Methods). However, no host could be predicted with a high degree of certainty.

Genomics analysis of phage Mar_Mega_1 identified a number of proteins detected in other megaphages including phage structural proteins and phage replication proteins (Table S2). Unlike some previously identified megaphages no CRISPR-cas system was identified [3]. However, a range of auxiliary metabolic genes (AMGs) were detected, homologues of which have not yet been identified in other megaphages (Table S2), including: nicotinamide-nucleotide amidohydrolase [PncC], isocitrate dehydrogenase [NADP] and a putative retropepsin-domain containing acid protease. Other AMGs present in Mar_Mega_1 included a patatin-like phospholipase (PLP) [11] which, together with a putative TonB-dependent receptor SusC were subsequently identified in LR745206, as well as AMGs encoding dihydrofolate reductase, a putative phosphoesterase and a putative peptidase which were also found in megaphages LR745206, LR756501, LR756502, LR756503 and LR756504 (Table S2).

The presence of AMGs potentially involved in carbon metabolism in Mar_Mega_1 is consistent with previous research indicating the prevalence of AMGs responsible for carbohydrate and amino acid uptake and metabolism in model marine phage systems and viral metagenomes [12]. For example, the TonB-dependent receptor SusC might be responsible for increasing carbohydrate uptake during infection [13], whereas isocitrate dehydrogenase [NADP], an AMG which was previously detected in marine viromes [14] carries out the oxidative decarboxylation of isocitrate to *α*-ketoglutarate (AKG). The latter product, AKG, is a rate-determining intermediate in the tricarboxylic acid cycle (EC 1.1.1.42) and crucial for both cellular energy metabolism and as a source of glutamate and glutamine and hence potentially important in both carbon and nitrogen metabolism during the infection process.

Furthermore, AMGs responsible for pyridine nucleotide synthesis such as nicotinamide-nucleotide amidohydrolase PncC (EC:3.5.1.42) whilst new to megaphages, have previously been found in other phages such as *Vibrio* phage KVP40 which encodes its own NAD^+^ salvage pathway [15]. Moreover, the Mar_Mega_1 phage encoded dihydrofolate reductase could act two-fold by increasing the host’s capacity to convert dihydrofolate into tetrahydrofolate (EC 1.5.1.3) which is essential for purine nucleotide biosynthesis or, due to similarity (identity 43%, coverage 100%) with a putative *dfrA3* antibiotic resistance gene, confer protection against diaminopyrimidine antibiotics, which are one of the most common antibiotic pollutants in marine environments [16].

This is the first time PLP was identified within a phage genome. Although the function of PLPs is currently not clear, a role in bacterial pathogen-eukaryotic host interactions was suggested [17]. We have since been able to identify a homologue of PLP in other phages (acc: LR745206), suggesting that megaphages might increase the virulence of their putative bacterial hosts.

## CONCLUSIONS

The complete genome of an abundant marine megaphage is described, which represents the largest phage genome documented to date from this environment. Using phylogenetic and genomic analyses it is distantly related to other megaphages and, based on current standards, represents a new family of phages.

## Materials and Methods

### Samples: collection and DNA extraction

Three marine water samples were collected (16.04.2019) from: Western Channel Observatory station L4 (50°15′00.0”N 4°13′01.2”W) surface waters, station L4 at the depth of 25 metres and Plymouth Sound (50°21′44.4”N 4°09′55.2”W) surface water. Water samples were filtered using 0.22 μm pore-size cellulose acetate membrane filters (Corning) to remove bacterial cells and larger particles. As megaphages have not been cultured their virion size is unknown, the larger genome size might result in larger capsids which might be excluded by a 0.22 μm pore-size filter. Subsequently, Amicon Ultra-15 100K (100,000 MWCO) concentrator columns (Millipore) were used to concentrate 4.5 L of each water sample, followed by 30 mL of SM wash buffer to remove the excess of marine salts. Resultant 1 mL of the permeate containing concentrated and cleaned up marine viral fraction in SM buffer was stored at 4°C. Non-encapsidated nucleic acids were removed by mixing 900 μL of each sample with 100 μL 10x DNase buffer and supplemented with 2 μL (4 U) of DNase I (NEB) and 1 U of RNase, followed by 30 min incubation at 37°C. Phage DNA was extracted using the phenol:chloroform:isoamyl alcohol [1], and stored at −20°C for further analysis.

### Short read sequencing (Illumina HiSeq)

Short read sequencing of the DNA samples was performed by the University of Liverpool Centre for Genomic Research using the Illumina HiSeq4000 platform (2 × 150 bp). Subsequently, Cutadapt v.1.2.1 (Martin, 2011) with ‘−O 3’ option was used to remove Illumina adapters from the resultant Fastq, followed by trimming using Sickle v.1.200 [2] with default options. Metagenomic contigs were assembled de novo using a previously described pipeline (Roux *et al*., 2019), as follows: tadpole.sh v.37.54 [3], using ‘mode=correct ecc=t prefilter=2’ was used for reads correction, followed by clumpify.sh v.37.54 [3] using ‘dedupe subs=0 passes=2’ options to remove the identical reads. Finally, concatenated reads from all three samples were cross-assembled using SPAdes v.3.13.0 [4] with ‘--sc -- only-assembler-k 77,99,127’ options.

### Long read sequencing (MinION)

Due to the low amount of available DNA and high requirements of the long-read sequencing, an aliquot of each DNA sample underwent whole genome amplification with the Illustra Ready-To-Go GenomiPhi V3 DNA Amplification Kit (GE Healthcare) following the manufacturer’s protocol. S1 Nuclease (Thermo Fisher Scientific) digestion was subsequently performed to de-branch chimeric DNA formed during the rolling circle amplification process ([5]). Resultant dsDNA was cleaned up using a DNA Clean & Concentrator Kit (Zymo Research) and used as an input for MinION sequencing.

MinION libraries were prepared using EXP-NBD104 and SQK-LSK109 kits (Oxford Nanopore) following the manufacturer’s protocol for the Native barcoding genomic DNA. Libraries were sequenced using MinION with a FLO-MIN106 flowcell and MinIT (Oxford Nanopore) for data storage and analysis. Bases were called with guppy_basecaller v.3.0.3 software using dna_r9.4.1_450bps_hac.cfg configuration for high accuracy calling, followed by demultiplexing of the samples with guppy_barcoder v.3.0.3 using default configuration and trimming of the barcodes with Porechop v.0.2.1 [6]. Genome assembly was performed using Flye v.2.6 [7] using ‘--nano-raw --meta --genome-size 5g --t 90’ options. Subsequently, minimap2 v.2.14 [8] with ‘-ax map-ont’ options was used to map MinION reads to the assembly.fasta file, followed by polishing of the output with marginPolish v.1.0.0 (Paten and Pesout, 2019) using ‘allParams.np.ecoli.json’ option. Finally, resultant assembly was further polished with Illumina short reads using Pilon v.1.23 (Walker *et al*., 2014) using the default options.

### Analysis of assembled metagenomes

Both Illumina and MinION metagenomics assemblies were concatenated and duplicate sequences and sub-sequences at >95% identity cut-off were removed with dedupe.sh [3] using ‘minidentity=95’ option. This was followed by the removal contigs shorter than 5,000 bp, which are unlikely to represent complete phage sequences. Remaining contigs were scanned with DeepVirFinder v.1.0 [9] using the default options with p value cutoff of <0.05 for the presence of phage sequences, followed by the checkV v.0.6 [10] analysis to determine the quality of the reconstructed phage genomes using: contamination, completeness, repeats and quality_summary commands with the default options.

Reconstructed megaphage gene sequences were predicted and annotated with Prokka v.1.11 [11] using a custom database of all phage genomes [12] that had previously been extracted from GenBank (Jul 2017) [12]. The resultant protein sequences were subsequently combined with protein sequences of all known phages and megaphages (Table S3) at the time of the analysis (January 2021) and used as an input for vContact2 v.0.9.15 [13] analysis with ‘--rel-mode Diamond --db ProkaryoticViralRefSeq94-Merged --pcs-mode MCL --vcs-mode ClusterONE’ options. Network map was visualised using Cytoscape v.3.7.2 [14]. Phylogenetic analysis of selected phages and all megaphages was based on the amino-acid sequence of terminase large subunit. Protein sequences were aligned using MAFFT v7.271 [15] with the default options followed by phylogenetic tree construction using IQ-TREE v.1.6.3 [16] using ‘--bb 1000’ option, with models of evolution selected using modeltest [17]. Resultant tree was visualised using ITOL [18] ANI was determined using Kostas lab online ANI calculator [19]. The closest known relatives vOTUs were identified by taking the top hit, when compared to a MASH database of current phage genomes [12]. The following mash settings were used: ‘mash -dist -d 0.3 -s 1000 -p 90’ [20]. Putative hosts of all vOTUs were predicted by comparison of vOTU contigs to a database of known CRISPR-spacers, allowing up to 3 mismatches in spacers [21].

Manual annotation of genes in Mar_mega_1 was carried out using Phyre2 [22] and HHpred [23] to predict the function of the large number of hypothetical proteins. Genomic comparison of Mar_Mega_1 with its two closest uncultured relatives (accession LR756502 and LR756502) [24] was carried out with get_homologues, with the following settings -M -t 0 -A [25].

### Viral abundance analysis

Viral abundance in both samples collected in this study and TARA dataset was determined by read mapping. Only short reads were used for estimation of abundance, to avoid bias introduced by amplification required for long read sequencing. Reads were mapped with bbmap.sh within minid=95 [3]. Viral abundance was determined using CPM (counts per million) as follows. For each contig with coverage >=70% and average fold >1.0x its number of reads was divided by contig’s length in kb resulting in reads per kilobase (RPK) value. CPM values were subsequently determined by dividing the RPK of each contig by the sum of all RPK values in the sample divided by a million. This value represented the abundance of each reconstructed phage genome normalised to the genome length and the total amount of reads obtained for each sample.

---

FigS1. Network map of Mar_Mega_1 in relation to known phages. Node colours denote: reference (NCBI) phage genomes (teal), Lak phages (yellow), megaphages (green) and Mar_Mega_1 (red).

TableS1. List of metagenomic contigs reconstructed in this study. Dataset includes: contig names, lengths, CheckV quality and completeness analysis, abundances (CPM) and closest known relative phage determination (where applicable).

TableS2. Functional annotation of the Mar_Mega_1 genome using Prokka and Phyre2 approaches. Dataset includes BlastP analysis and homologue search results for selected putative auxiliary metabolic genes.

TableS3. List of megaphages and Lak phages (including incomplete megaphage genomes) used in vContact2 and phylogenetic analyses.

## References

1. Dion MB, Oechslin F, Moineau S. Phage diversity, genomics and phylogeny. Nat Rev Microbiol 2020; 18:125–138.

2. Yuan Y, Gao M. Jumbo bacteriophages: An overview. Front Microbiol 2017.

3. Devoto AE, Santini JM, Olm MR, Anantharaman K, Munk P, Tung J, et al. Megaphages infect *Prevotella* and variants are widespread in gut microbiomes. Nature Microbiology 2019.

4. Al-Shayeb B, Sachdeva R, Chen L-X, Ward F, Munk P, Devoto A, et al. Clades of huge phages from across Earth’s ecosystems. Nature 2020; 578: 425–431.

5. Crisci MA, Chen LX, Devoto AE, Borges AL, Bordin N. Wide distribution of alternatively coded Lak megaphages in animal microbiomes. bioRxiv 2021.

6. Chen L-X, Méheust R, Crits-Christoph A, McMahon KD, Nelson TC, Slater GF, et al. Large freshwater phages with the potential to augment aerobic methane oxidation. Nat Microbiol 2020; 5:1504–1515.

7. Roux S, Adriaenssens EM, Dutilh BE, Koonin EV, Kropinski AM, Krupovic M, et al. Minimum information about an uncultivated virus genome (MIUViG). Nat Biotechnol 2019; 37: 29–37.

8. Turner D, Kropinski AM, Adriaenssens EM. A roadmap for genome-based phage taxonomy. Viruses 2021; 13.

9. Brum JR, Ignacio-espinoza JC, Roux S, Doulcier G, Acinas SG, Alberti A, et al. Ocean Viral Communities. Science 2015; 348: 1261498–1–11.

10. Roux S, Emerson JB, Eloe-Fadrosh EA, Sullivan MB. Benchmarking viromics: an in silico evaluation of metagenome-enabled estimates of viral community composition and diversity. PeerJ 2017; 5: e3817.

11. Wilson SK, Knoll LJ. Patatin-like phospholipases in microbial infections with emerging roles in fatty acid metabolism and immune regulation by Apicomplexa. Mol Microbiol 2018; 107: 34–46.

12. Warwick-dugdale J, Buchholz HH, Allen MJ, Temperton B. Host-hijacking and planktonic piracy : how phages command the microbial high seas. Virology J 2019; 1: 1–13.

13. Reeves AR, D’Elia JN, Frias J, Salyers AA. A Bacteroides thetaiotaomicron outer membrane protein that is essential for utilization of maltooligosaccharides and starch. J Bacteriol 1996; 178:823–830.

14. Hurwitz BL, Hallam SJ, Sullivan MB. Metabolic reprogramming by viruses in the sunlit and dark ocean. Genome Biol 2013; 14: R123.

15. Lee JY, Li Z, Miller ES. Vibrio Phage KVP40 encodes a functional NAD+ salvage pathway. J Bacteriol 2017; 199.

16. Felis E, Kalka J, Sochacki A, Kowalska K, Bajkacz S, Harnisz M, et al. Antimicrobial pharmaceuticals in the aquatic environment - occurrence and environmental implications. Eur J Pharmacol 2020; 866: 172813.

17. Banerji S, Flieger A. Patatin-like proteins: a new family of lipolytic enzymes present in bacteria? Microbiology 2004; 150: 522–525.

## References

1. Cook R, Hooton S, Trivedi U, King L, Dodd CER, Hobman JL, et al. Hybrid assembly of an agricultural slurry virome reveals a diverse and stable community with the potential to alter the metabolism and virulence of veterinary pathogens. Microbiome 2021; 9: 65.

2. Joshi NA, Fass JN, Others. Sickle: A sliding-window, adaptive, quality-based trimming tool for FastQ files (Version 1.33)[Software]. 2011.

3. Bushnell B. BBMap: A fast, accurate, splice-aware aligner. 2014. Lawrence Berkeley National Lab. (LBNL), Berkeley, CA (United States).

4. Bankevich A, Nurk S, Antipov D, Gurevich A a., Dvorkin M, Kulikov AS, et al. SPAdes: A New Genome Assembly Algorithm and Its Applications to Single-Cell Sequencing. J Comput Biol 2012; 19:455–477.

5. Lasken RS, Stockwell TB. Mechanism of chimera formation during the Multiple Displacement Amplification reaction. BMC Biotechnol 2007; 7: 1–11.

6. Wick RR. Porechop. Github https://github.com/rrwick. 2017. Porechop.

7. Kolmogorov M, Bickhart DM, Behsaz B, Gurevich A, Rayko M, Shin SB, et al. metaFlye: scalable long-read metagenome assembly using repeat graphs. Nat Methods 2020; 17: 1103–1110.

8. Li H. Minimap2: pairwise alignment for nucleotide sequences. Bioinformatics 2018; 34: 3094–3100.

9. Ren J, Song K, Deng C, Ahlgren NA, Fuhrman JA, Li Y, et al. Identifying viruses from metagenomic data by deep learning. 2018.

10. Nayfach S, Camargo AP, Schulz F, Eloe-Fadrosh E, Roux S, Kyrpides NC. CheckV assesses the quality and completeness of metagenome-assembled viral genomes. Nat Biotechnol 2021; 39: 578–585.

11. Seemann T. Prokka: Rapid prokaryotic genome annotation. Bioinformatics 2014; 30: 2068–2069.

12. Cook R, Brown N, Redgwell T, Rihtman B, Barnes M, Clokie M, et al. INfrastructure for a PHAge REference Database: Identification of large-scale biases in the current collection of phage genomes. bioRxiv. 2021., 2021.05.01.442102

13. Bin Jang H, Bolduc B, Zablocki O, Kuhn JH, Roux S, Adriaenssens EM, et al. Taxonomic assignment of uncultivated prokaryotic virus genomes is enabled by gene-sharing networks. Nat Biotechnol 2019.

14. Shannon P, Markiel A, Ozier O, Baliga NS, Wang JT, Ramage D, et al. Cytoscape: a software environment for integrated models of biomolecular interaction networks. Genome Res 2003; 13:2498–2504.

15. Katoh K, Standley DM. MAFFT multiple sequence alignment software version 7: improvements in performance and usability. Mol Biol Evol 2013; 30: 772–780.

16. Nguyen L-T, Schmidt HA, von Haeseler A, Minh BQ. IQ-TREE: A fast and effective stochastic algorithm for estimating maximum-likelihood phylogenies. Mol Biol Evol 2015; 32: 268–274.

17. Posada D, Crandall KA. MODELTEST: testing the model of DNA substitution. Bioinformatics 1998; 14:817–818.

18. Letunic I, Bork P. Interactive Tree Of Life (iTOL): An online tool for phylogenetic tree display and annotation. Bioinformatics 2007; 23: 127–128.

19. Rodriguez-R LM, Konstantinidis KT. Bypassing cultivation to identify bacterial species. Microbe Wash DC 2014; 9: 111–118.

20. Ondov BD, Treangen TJ, Melsted P, Mallonee AB, Bergman NH, Koren S, et al. Mash: fast genome and metagenome distance estimation using MinHash. Genome Biol 2016; 17: 132.

21. Dion MB, Plante P-L, Zufferey E, Shah SA, Corbeil J, Moineau S. Streamlining CRISPR spacer-based bacterial host predictions to decipher the viral dark matter. Nucleic Acids Res 2021; 49: 3127–3138.

22. Kelley LA, Mezulis S, Yates CM, Wass MN, Sternberg MJE. The Phyre2 web portal for protein modeling, prediction and analysis. Nat Protoc 2015; 10: 845–858.

23. Söding J, Biegert A, Lupas AN. The HHpred interactive server for protein homology detection and structure prediction. Nucleic Acids Res 2005; 33: W244–8.

24. Devoto AE, Santini JM, Olm MR, Anantharaman K, Munk P, Tung J, et al. Megaphages infect *Prevotella* and variants are widespread in gut microbiomes. Nature Microbiology 2019.

25. Contreras-Moreira B, Vinuesa P. GET_HOMOLOGUES, a versatile software package for scalable and robust microbial pangenome analysis. Appl Environ Microbiol 2013; 79: 7696–7701.

